# Identifying Phage Host Receptors Using TraDIS

**DOI:** 10.1101/2025.12.01.691526

**Authors:** Byron Gentle, Henry Scott, Claire Maher, Ian Grainge

## Abstract

The growing antimicrobial resistance crisis has sparked renewed interest in using bacteriophage (phage) as a treatment for antibiotic-resistant infections. Phage are viruses that infect and often kill bacteria, and have served as key models for many molecular biology studies. There are many complex phage-bacterial interactions that dictate the success or failure of the phage life cycle, but many of these are not well understood or characterised. These include the specific interactions to recognize a bacterium during the infection process, bacterial phage defence mechanisms and metabolic pathways that are key to phage replication. In this study a novel *Escherichia coli* phage, Arnold, is characterised and the genome sequence is reported. We then use transposon-directed insertion sequencing (TraDIS) to screen a library of *E. coli* mutants to identify over 200 genes involved in Arnold phage infection or in phage resistance. Using this approach, we identified both an outer membrane protein (BtuB, vitamin B12 transporter) and an inner membrane protein (SdaC, serine/proton symporter) involved in phage infection. Deletions in these genes were shown to provide immunity to infection by this phage. Understanding the genetic basis of phage infection and phage resistance should allow us to select or design phage to target specific bacterial pathogens in the future.

## Introduction

Since the development of antibiotics, their consistent misuse has facilitated the emergence and spread of antimicrobial resistance (AMR). Despite the increasing number and variety of AMR bacteria, the development of new antibiotics has declined significantly (Cooper & Shlaes, 2011). This sharp reduction in antibiotic discovery is concurrent with the evolution of multidrug-resistant or pan-drug resistant bacteria, which have developed mechanisms to resist all known antibiotics, resulting in incurable infections that often lead to death. The World Health Organisation (WHO) has identified AMR as one of the top 10 global health threats (World Health Organisation, 2021). As of 2016, AMR was attributed to approximately 700,000 deaths annually, with predictions of reaching 10 million deaths per year by 2050 (O’Neill, 2014). Alarmingly, the death toll from AMR has already surged towards that figure; by 2019 there were 4.95 million deaths associated with AMR, of which 1.2 million were directly attributable (Antimicrobial Resistance, 2022). If this trend continues, annual AMR-related deaths are projected to surpass the deaths from all cancers within 25 years. The recognition of AMR as a critical public health threat has driven demand for a deeper understanding of bacteria’s ability to evolve antibiotic resistance and the development of newer, more effective antimicrobial treatments, such as bacteriophage (phage) therapy.

Bacteriophage therapy has existed for over a century, with its discovery and first use in the early 1900s (Salmond & Fineran, 2015). Phage therapy makes use of bacteriophage to attack and kill bacteria as an alternative treatment to antibiotics. Despite phage therapy’s initial success, its use was sidelined (at least by Western medicine) with the discovery and success of antibiotics, which proved easier to prescribe and had broad spectrum uses. Conversely, phage are often very specific for certain species or even strains of bacteria. This specificity is largely driven by the specific recognition of receptors on the bacterial cell surface. However, due to the recent growing AMR crisis, phage therapy is being re-visited by the scientific and health communities as an alternative to antibiotics (Gilbey et al., 2019). As phage are the natural predators to bacteria, it is logical to assume that at least one phage exists against any given bacterial strain. This would allow for the potential isolation and development of phage therapy against any bacterial infection necessary.

Host recognition is a vital part of a bacteriophage’s life cycle, allowing for the selection of the phage’s prey species and subsequent infection. This occurs when a phage identifies and binds to a component on a bacterial cell surface; this receptor is specific and generally consists of either an outer-membrane protein, lipopolysaccharides, pili, capsule or flagellar component (Letellier et al., 1999). This receptor binding is followed by the release of the phage’s genetic material into the host cell where it can be transcribed/translated and replicated to produce more phage. To enable the release of genetic material into the cytoplasm the outer leaflet of the bacterial cell must be penetrated and mechanisms to achieve this vary greatly between phage.

Here we report the sequence of an *Escherichia coli*-specific phage, Arnold, which has a 50,935 bp genome. It was identified in a screen for phage that could infect two separate *E. coli* K12 lab strains- BW25113 and AB1157. Transposon-directed insertion-site sequencing (TraDIS), also called TnSeq, was used to identify genes involved in phage infection and phage resistance. A high-density transposon mutant library of BW25113 (Alquethamy et al., 2021, Maher et al., 2022) was challenged with Arnold phage and the profile of surviving mutant cells was compared to a non-infected control. 222 significantly over or under-represented genes were identified after phage challenge. Mutants of the genes that were the top 4 most significantly overrepresented and top 2 underrepresented hits from this screen were challenged with Arnold phage. Two of these mutants were shown to be resistant to phage infection, identifying them as putative phage attachment sites: BtuB, an outer membrane transporter, and SdaC, an inner membrane transporter. These results suggest Arnold could be a close relative of the C1 phage identified and studied in Russia in the 1990s that uses the same inner and outer membrane receptor proteins (Likhacheva et al., 1990; Likhacheva et al., 1996).

## Materials and Methods

### Arnold isolation and sequencing

Arnold was isolated in a screen for phage that could infect two *E. coli* K12 lab strains, AB1157 and BW25113. DNA was isolated and sequenced using Illumina sequencing (SeqCentre, USA). Trimmed paired-end reads were used to assemble the genome using Shovill (Seeman, 2020). To determine quality, raw reads were aligned against the phage DNA contig using Bowtie2 (Langmead & Salzberg, 2012). Gene identification and annotation was carried out using Pharokka (Bouras et al., 2022).

### TraDIS challenge and sequencing

A library of random Tn5 insertion mutants had been generated (Maher et al., 2022) with an estimated 400,000 unique insertion mutants. The mutant library was grown until ∼ 10^9^ bacterial cells were present, and these were infected with 10^10^ phage. The mix was transferred to 10 ml fresh LB medium and grown at 37°C for 5 hours with agitation. A control was also made to the same volume with 10^9^ bacterial cells from the TraDIS library, however, no phage was added. Both the control and challenge were done in biological duplicates. Following incubation, the cultures were centrifuged at 6000xg for 10 minutes at 4°C. The supernatant was then discarded, and the pellet was resuspended in 10 mM Tris (pH 7.5). DNA extraction was conducted using the Qiagen DNeasy UltraClean Microbial Kit as per the manufacturer’s protocol. The resulting DNA quality and concentration was confirmed using a NanoDrop spectrophotometer (NanoDrop® ND-1000 UV-Vis Spectrophotometer, ThermoFisher) and on a Qubit (Qubit™ 4 Flurometer, ThermoFisher). DNA was sent for TraDIS sequencing at UNSW’s Ramaciotti Centre for Genomics, using established protocols (Alquethamy et al., 2021). The resulting reads from the sequencing were then analysed following the Bio-TraDIS analysis pipeline (Barquist et al, 2016).

### *BtuB* deletion

Deletion was carried out using Lambda Red recombineering as previously described (Datsenko & Wanner, 2000). A kanamycin resistance gene cassette bearing homology to the *E. coli btuB* gene at either end was amplified by two steps of PCR. First, the *btuB* homology regions were amplified from BW25113 genomic DNA using the following primers: *btuB* upstream for 5′ GCGAATCTGGAGCTGAC 3′, *btuB* upstream rev 5′ GATATCAAGCTTATCGATACCGTCTTATCACATTGTAAAGCATCCAC 3′ and *btuB* internal for 5′ ACTAGTTCTAGAGCGGCCGCGGCAAACCAGCGCCG 3′, *btuB* internal rev 5′ CAAGTCACTGACATCGTTACG 3′. The resulting products were then used to amplify the kanamycin resistance gene from the plasmid pBluescript::FRT-Kan-FRT. This PCR product served as the kanamycin cassette with homology to sites within the *btuB* gene.

BW25113 was transformed with the plasmid pKD46 and plated onto LB agar containing 100 µg/ml ampicillin. Subsequently these cells were then transformed with the linear PCR product containing the kanamycin resistance gene cassette, and selected on LB agar containing 50 µg/ml kanamycin. Colonies were screened using the *btuB* upstream for primer (above) and a screening primer 5′ CTTCGATGAATTCCCAACCG 3′ to amplify the cassette at a length of 1325bp. PCR products were sequenced to confirm the correct insertion.

### Growth and phage challenge of selected BW25113 mutants

The *sdaC, yfdF, ybeT*, *ygfA* and *moaD* mutants were revived from the Keio collection and grown in LB with 25μg/ml kanamycin. Each strain, plus the *btuB* deletion and the wt control was grown to an OD of 0.2 and then 10 μl was added to 180 μl of fresh LB in a 96 well plate. Six wells were inoculated for each strain, and to three of these phage Arnold was added at a MOI of 1, with the other three being uninfected controls, with PBS added instead of phage. Plates were then placed in a LogPhase600 (BioTek) and grown at 37°C with 300rpm shaking between readings. OD readings were taken every 5 minutes.

## Results

### Phage Arnold Isolation

The phage investigated in this study was detected during experiments aimed at broadening phage host range. Subsequent analysis revealed that it was not a product of directed host-range training but instead represented a contaminant of unknown origin. It was found to be able to infect two different *E. coli* K12 lab strains, AB1157 and BW25113. This phage, named Arnold, was purified by multiple rounds of plaque isolation using the double-agar overlay method and was subsequently characterised through sequencing and genome assembly. Illumina sequencing of Arnold generated a total of 4.5 million paired end reads. From these reads a single contig of 50,935 bp was assembled, and 99.22% of the reads mapped to this contig. When the genome was analysed in Pharokka (Bouras et al., 2022), a total of 100 genes were identified. From these 100 genes, 36 were identified as having a recognisable homologue and the remaining 64 were left as hypothetical genes (Figure 1). A BLASTn search using phage Arnold’s genome showed the closest match was *Escherichia* phage vB_EcoS-G3B1, being 97% identical at the DNA level. This phage belongs to the Caudoviricetes class and Warwickvirus genus.

**Figure 1:**
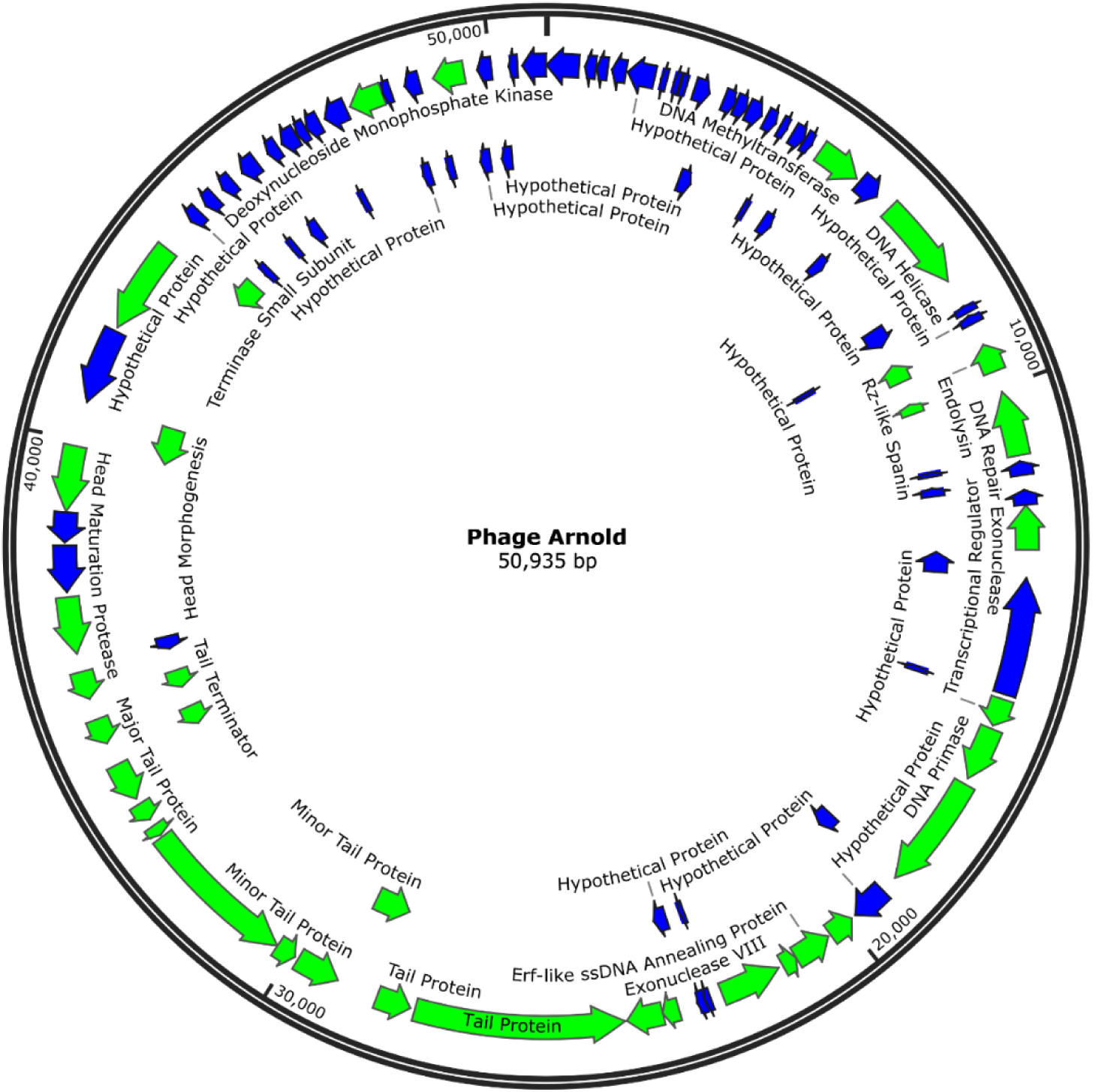
Circularised representation of the Phage Arnold genome. Hypothetical proteins are in blue while proteins with a predicted function (Pharokka) are in green.

### TraDIS Analysis

To identify host genes involved in phage susceptibility and in phage resistance, TraDIS (or TnSeq) was used. A library of random Tn5 insertion mutants had been generated (Maher et al., 2022) with an estimated 400,000 unique insertion mutants in the strain BW25113. The mutant library was grown until ∼ 10^9^ bacterial cells were present, and these were infected at a high multiplicity of infection (MOI = 10). After 5 hours of growth the surviving cells were collected and DNA was extracted and sequenced.

The resulting reads were processed through the bioTraDIS pipeline to map the insertion sites in both control and challenge samples (Barquist et al., 2016), and the log fold change (LFC) of insertions after phage challenge for each gene was calculated. A positive LFC indicates that the inserted transposon/ disrupted gene conferred a survival advantage for the bacteria in the presence of the phage, whereas a negative LFC suggests that the phage more readily killed that mutant compared to the wild type. The LFC was plotted against the calculated statistical significance of the change in insertions in each gene, represented by adjusted P values (Q values) (Figure 2). A total of 222 genes were identified as statistically significant, two of which received such a low Q value it registered as a zero. A full table of all 222 genes is shown in Supplementary Table 1.

**Figure 2.**
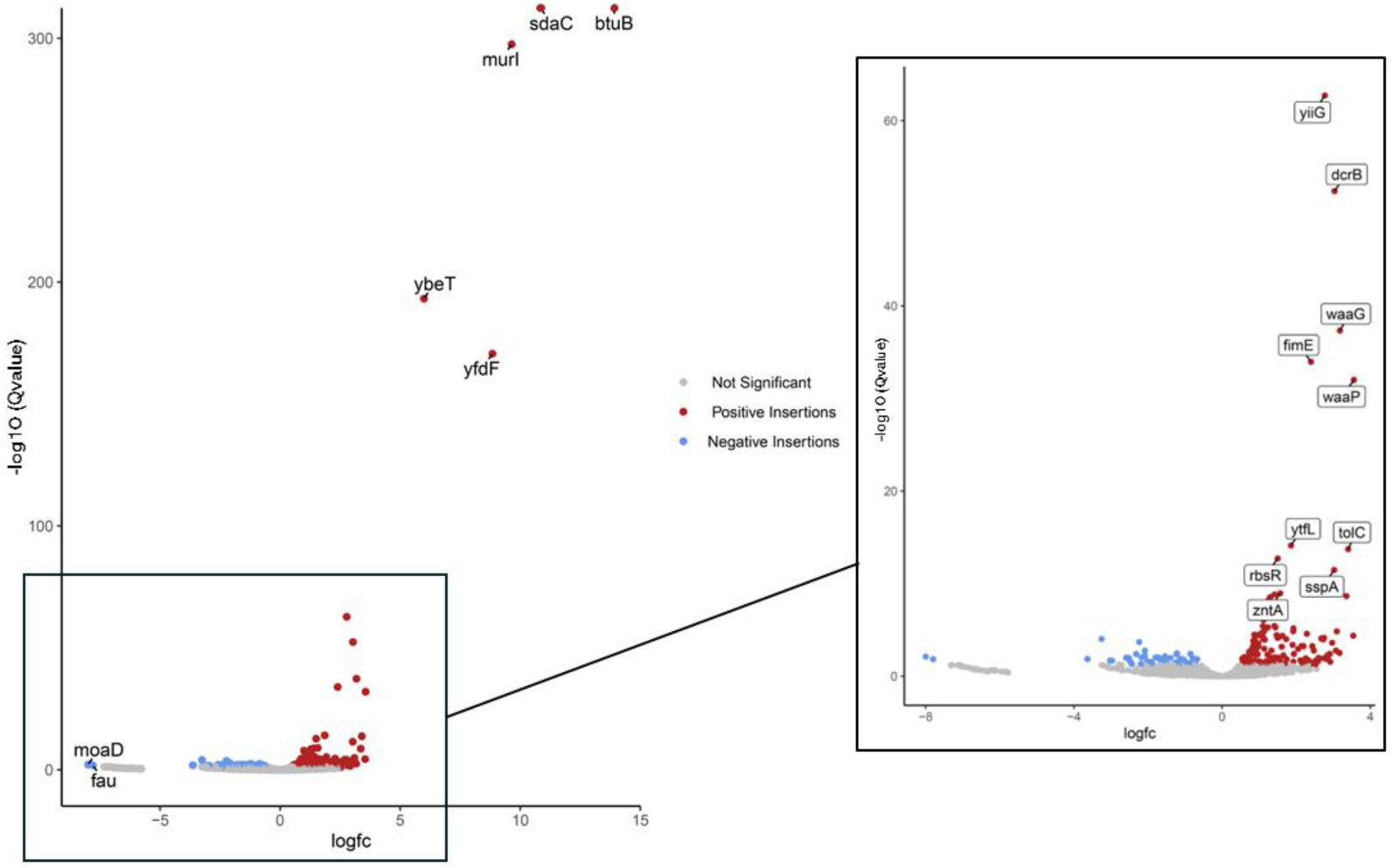
Volcano plot of Log Fold Change (Log_2_Fc) vs -log_10_(Q value) for the TraDIS sequencing. Insertions with a Q value significance of <0.05 are highlighted in red for a positive LogFc and blue for a negative LogFc. The 5 most significant positive hits are labelled along with the two most significant negative hits. On the right is shown an expanded view of the boxed section with the next ten most significant genes annotated.

The two genes with the greatest increase in transposon insertions had Q values approaching zero: *btuB* (Vitamin B12 outer membrane transporter) (Log_2_FC = 13.9 and Q <10^-300^) and *sdaC* (L-serine inner membrane proton symporter) (Log_2_FC = 10.8 and Q <10^-300^). These were followed in significance by *murI* (cytosol glutamate racemase), and two uncharacterised genes *ybeT* and *yfdF*. These top five most significant genes were investigated further to identify their potential role in phage infection. Additionally, the two genes showing the largest negative LFC, *moaD* (molybdopterin synthase Sulphur carrier subunit) (Log_2_FC = -8.0 and Q = 0.0076) and *ygfA*, also called *fau*, (putative 5-formyltetrahydrofolate cyclo-ligase) (Log_2_FC = -7.9 and Q = 0.014)were also chosen for further study. Although significant, their Q values were not as remarkable as those of the top positive insertions.

One of the top 5 genes with the greatest increase in insertions after Arnold infection, *murI,* has partial overlap with the end of the *btuB* gene. Insertions in *murI* were only identified in this overlap region, while the rest of the gene sequence showed no insertions. This suggests that identification of *murI* is likely an artefact due to its shared genetic sequence with *btuB* and that *murI* itself may not impact the phage’s ability to infect the host. In fact, *murI* is essential in *E. coli* and cannot be deleted. The presence of some insertions in the 5’ portion of the gene that overlaps with *btuB* is perhaps surprising then, as it presumably must somehow still produce an active protein. There may be a possible alternate start site that can be used that may remove the very N-terminus of the protein but retain function. Insertions of transposons in essential genes, often in the 3’ end or other non-essential regions, are not uncommon in TraDIS data sets (Christen et al., 2011; Goodall et al., 2018). Further, the insertions in the *btuB/murI* overlapping portion are highly polarised with roughly 10-fold higher insertions rates in one direction compared to the other (Supplementary figure 1). It has been shown that the Tn5 transposon can provide transcription into a neighbouring gene if in the correct orientation, as well as alternate translation initiation sites (Goodall et al., 2018). For this reason *murI* was not investigated further for its role in phage infection.

12 genes involved in core LPS biosynthesis were also identified using TraDIS: *waaA*, *waaB*, *waaG*, *waaJ*, *waaL*, *waaP*, *waaQ*, *waaR*, *waaS*, *waaU*, *waaY* and *waaZ*. These all had positive Log_2_FC values between 3.56 and 0.74, with two genes, *waaP* and *waaG*, being in the top ten largest fold-change hits of the screen. In addition, another five genes with LPS modifying or regulating functions were identified: *wzzE, wecH, yejM, arnF* and *rfaH*. This strongly suggests a role for LPS in aiding the infection of the phage Arnold, and interaction with LPS during initial attachment has been observed for other phage like T4 and T7 (Lindberg, 1973; Washizaki et al., 2016). However, this was not investigated further here, but is an avenue for future study.

### Confirmation of TraDIS results using gene knockouts

Individual knockouts of the top 6 identified genes from TraDIS (4 positive LFC and 2 negative LFC) were used to investigate phage Arnold infection kinetics. The single knockouts of *sdaC, yfdF, ybeT, ygfA* and *moaD* were obtained from the Keio library. The Keio collection of *E. coli* BW25113 does not contain a knockout of the *btuB* gene, due to the gene sequence partially overlapping the essential gene *murI*. Neither gene could be deleted in the original Keio screen (Baba et al., 2006). Lambda Red was utilised to recombine a kanamycin resistance cassette into the *btuB* gene, deleting a large portion of the gene (the first ∼1000 bp including the ATG start codon was removed) but leaving the neighbouring *murI* intact. The subsequent knockout was viable and the deletion of the majority of *btuB* was confirmed by both PCR and DNA sequencing.

BW25113 and the single gene knockouts were grown to an OD_600_ of ∼0.2. Aliquots of cells were added to a 96-well plate with fresh media and infected with Arnold phage at a MOI of 1 in triplicate. Plates were incubated in a plate reader shaking at 300 rpm at 37°C and the optical density was read every five minutes over a total of 70 minutes (Figure x). Both the *btuB* and *sdaC* knockouts showed no detectable inhibition of growth or cell death from the phage infection during the assay. These results point to a key role in infection played by BtuB and SdaC, which are outer and inner membrane proteins respectively. The most likely role is that they are the receptors to which the phage binds or play a key part in injection of the phage genome. These two genes were identified by genetic studies as the likely receptors for the *E. coli* K12 lytic phage C1 (Letellier et al., 1999; Likhacheva et al., 1996); inactivating mutants in each gene were found to prevent C1 adsorption and infection. In addition C1 phage was seen to be dependent upon the DcrB protein, a periplasmic protein. *dcrB* was found in the TraDIS sequencing as a significant hit after Arnold challenge (Log_2_FC = 3.0, Q value = 4 x 10^-53^). This strongly suggests that Arnold is closely related to C1 phage as it relies on the same 3 genes for infection, although we were unable to find a deposited genome sequence for C1 to confirm this.

The knockouts for *yfdF, ybeT*, *ygfA* and *moaD* all displayed similar growth and subsequent cell death following infection as the wild-type control strain within the sensitivity of this assay.

## Discussion

We have identified a phage, Arnold, that targets two *E. coli* K12 strains, BW25113 and AB1157. It has some close relatives (97% identity at the DNA level) whose genomes have been deposited, but we believe it is also related to the previously studied C1-like phages. Using TraDIS, over 200 genes were identified as linked to either susceptibility or resistance to the phage. Two genes, the outer membrane transporter BtuB and inner membrane serine-proton symporter SdaC, were found to be essential for infection. In addition, multiple genes involved in lipopolysaccharide biosynthesis were identified as being important for infection. This work provides insight into the genetic requirement for Arnold and C1-like phage infection of the *E. coli* host, and highlights the value of transposon mutagenesis screening and subsequent sequencing for identification of phage receptors.

### BtuB as a phage receptor

The most significant hit in the TraDIS screen was the vitamin B12 transporter protein, BtuB. BtuB is a well-established receptor for multiple coliphages, including T5-like phages BF23, ϕR2-01 and GSP044, as well as phages infecting *Yersinia* and *Salmonella* (Cumby et al., 2015; Letellier et al., 1999). We constructed a partial deletion of *btuB,* which completely abolished Arnold infection, confirming that this is likely to be the primary phage attachment site (Figure 4). The 3’ end of the *btuB* gene overlaps with the 5’ end of the essential *murI* gene meaning that a complete knockout could not be generated, and was absent from the Keio collection. Interestingly, the presence of transposon insertions in the 5′ region of *murI* that overlaps *btuB* suggests that MurI can still function when its N-terminus is somewhat truncated, perhaps by use of an alternative start codon.

**Figure 3.**
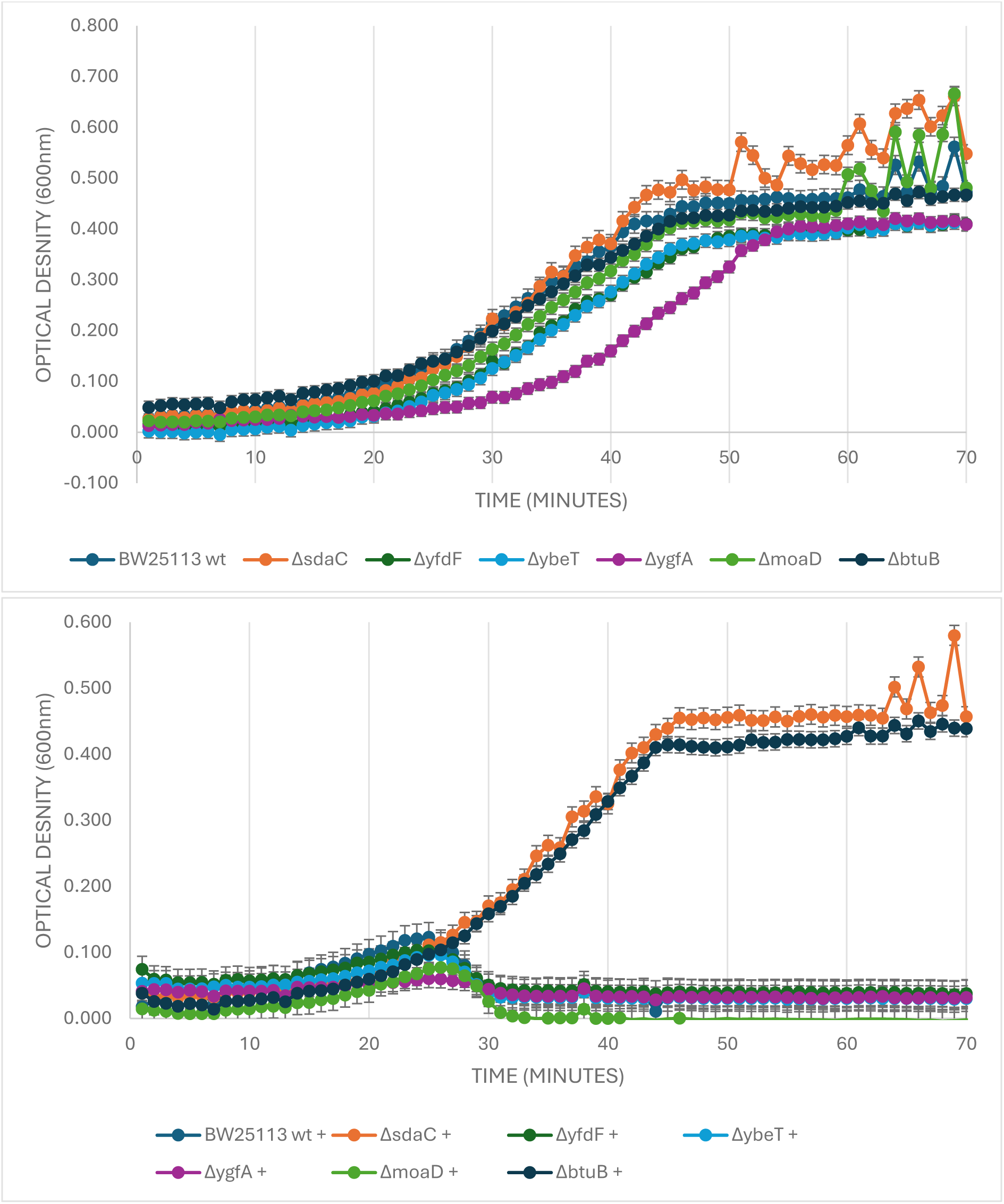
Optical density measurements of growth of *E. coli* BW25113 and selected mutants over time at 37°C, with and without phage Arnold infection. (A) OD readings at 600nm over 70 minutes of BW25113 and knockout mutants: *sdaC, yfdF, ybeT, ygfA, moaD*, and *btuB*. (B) OD readings for the same strains with phage Arnold infection at an MOI of 1 at time zero.

**Figure 4:**
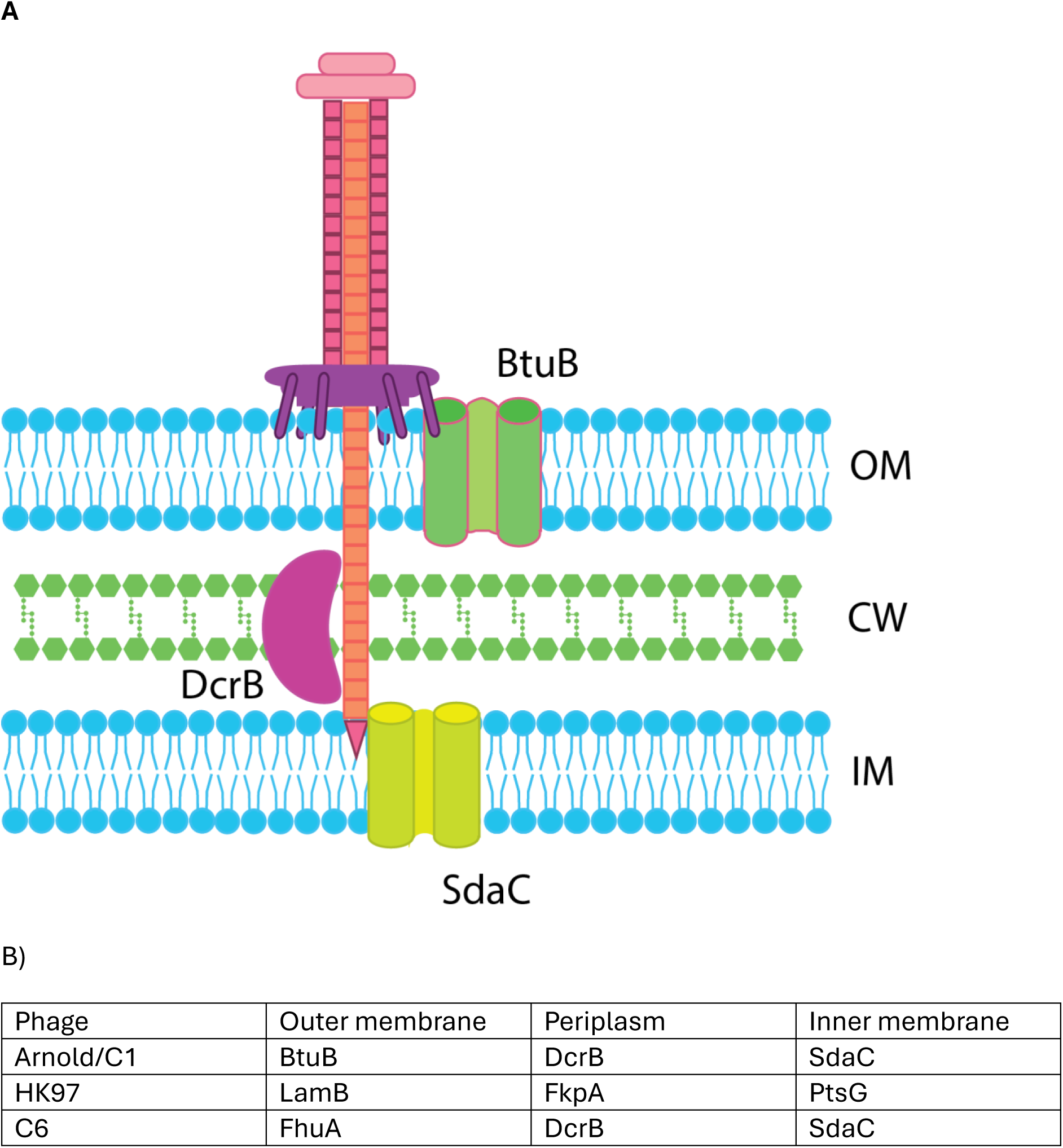
A) Cartoon representation of Arnold infection. The model suggest Arnold contacts the outer membrane (OM) transport protein BtuB and the inner membrane (IM) transport protein SdaC, as well as interacting with the periplasmic protein DcrB. CW= cell wall. B) The table shows the different requirement for Arnold/C1, HK97 and C6 phage.

### SdaC as a secondary/inner membrane receptor

The second major TraDIS hit was the *sdaC* gene encoding an inner membrane L-serine/proton symporter. This gene was previously shown to be involved in infection by C1 and C6 phage, with the suggestion that it was involved in the DNA injection process, perhaps being used as a channel for DNA to cross the inner membrane (Likhacheva et al., 1996; Samsonov et al., 2002). The complete resistance of the *sdaC* knockout to Arnold infection suggests that SdaC is essential for a second-stage recognition or DNA injection step, fitting a sequential receptor model.

Phage C1 has also been shown to require the periplasmic DrcB protein, and the *dcrB* gene was also identified as a very significant hit in the TraDIS screen for Arnold (Table S1). In C6 phage, the outer membrane receptor is FhuA, but it also requires both SdaC and DcrB for successful infection, highlighting that the dual-receptor strategy involving inner membrane and periplasmic proteins are conserved. A similar strategy was observed with the HK97 phage (Cumby et al., 2015), where the phage attaches to the outer membrane transporter LamB, but also requires the periplasmic protein FkpA and inner membrane protein PtsG (Figure 4). In this case the phage tape-measure protein (TMP) of the phage was shown to interact with PtsG, and was thought to constitute the channel through the inner membrane whereby the phage DNA could enter the cell. Exactly how widespread the requirement for periplasmic and inner-membrane proteins are for phage remains to be seen, as too few have been extensively characterised.

A multi-step or dual-receptor strategy may be used by phage to increase host specificity, but possibly at the expense of restricting their host range to only those cells expressing all the necessary components. Dual receptor proteins may also have co-evolved with super-infection exclusion proteins that inhibit passage of the phage DNA across the cell envelope. At least in some cases this has been shown to act following adsorption to the cell surface, preventing a channel from forming to allow DNA passage (Cumby et al., 2012).

### Role of LPS biosynthetic genes

In total 17 genes involved in the LPS biosynthesis pathway were significantly enriched in the TraDIS dataset, with two, *waaP* and *waaG,* being in the top ten positive fold-change hits. This enrichment may indicate a direct role for LPS in Arnold adsorption, either as a co-receptor or stabilising factor, or an indirect role in supporting correct folding and localisation of BtuB within the outer membrane. Further work would be needed to distinguish these possibilities. Similar dependencies on intact LPS have been reported for phages that otherwise target outer membrane proteins or flagellar proteins, as truncations in the LPS core can disrupt receptor accessibility or initial adherence to the cell (Cowley et al., 2018; Harding et al., 2025).

### Negative LogFoldChange genes

The two most significant genes with negative LFC values were *moaD*, encoding the molybdopterin synthase sulphur carrier subunit, and *ygfA* (also known as *fau*), a putative 5-formyltetrahydrofolate cyclo-ligase. Their disruption appeared to increase susceptibility to Arnold infection, suggesting that they may contribute to resistance under normal conditions. Although their precise roles remain unclear, altered metabolism could influence envelope composition or receptor expression, thereby enhancing infection efficiency. Alternatively, some of the genes with negative LogFC could influence biofilm formation, such as *bssS* (Log_2_FC ∼ -2.5) which is a positive regulator of biofilm formation (Domka et al., 2006). Further study is required to confirm these effects.

### Implications for phage therapy and resistance evolution

Identifying host receptors is crucial for the rational selection and engineering of phages for therapeutic applications. Knowledge of receptor proteins allows prediction of potential resistance mechanisms and facilitates the design of phage cocktails targeting multiple independent receptors. It is unclear if Arnold’s requirement for both BtuB and SdaC would make the evolution of resistance slower, as simultaneous mutations in both loci would be needed to confer complete immunity, or if it would enhance resistance as it gives two possible gene targets where mutations would abolish infection. The specificity shown by utilising a dual receptor strategy does seem to narrow the host range and possibly argues against usefulness against diverse clinical isolates lacking one or both receptors.

In conclusion, Arnold is a C1-like *E. coli* phage that requires both the outer membrane transporter BtuB and the inner membrane symporter SdaC for successful infection, with additional contributions from the periplasmic protein DcrB and intact LPS biosynthesis. This dual-receptor requirement mirrors that of several other phages and underscores the complexity of phage–host interactions. The combination of genome sequencing with TraDIS has provided a comprehensive view of the genetic determinants of Arnold infection, supporting future efforts to exploit phages in the fight against antimicrobial-resistant bacteria.

---

**Supplementary Figure 1:**
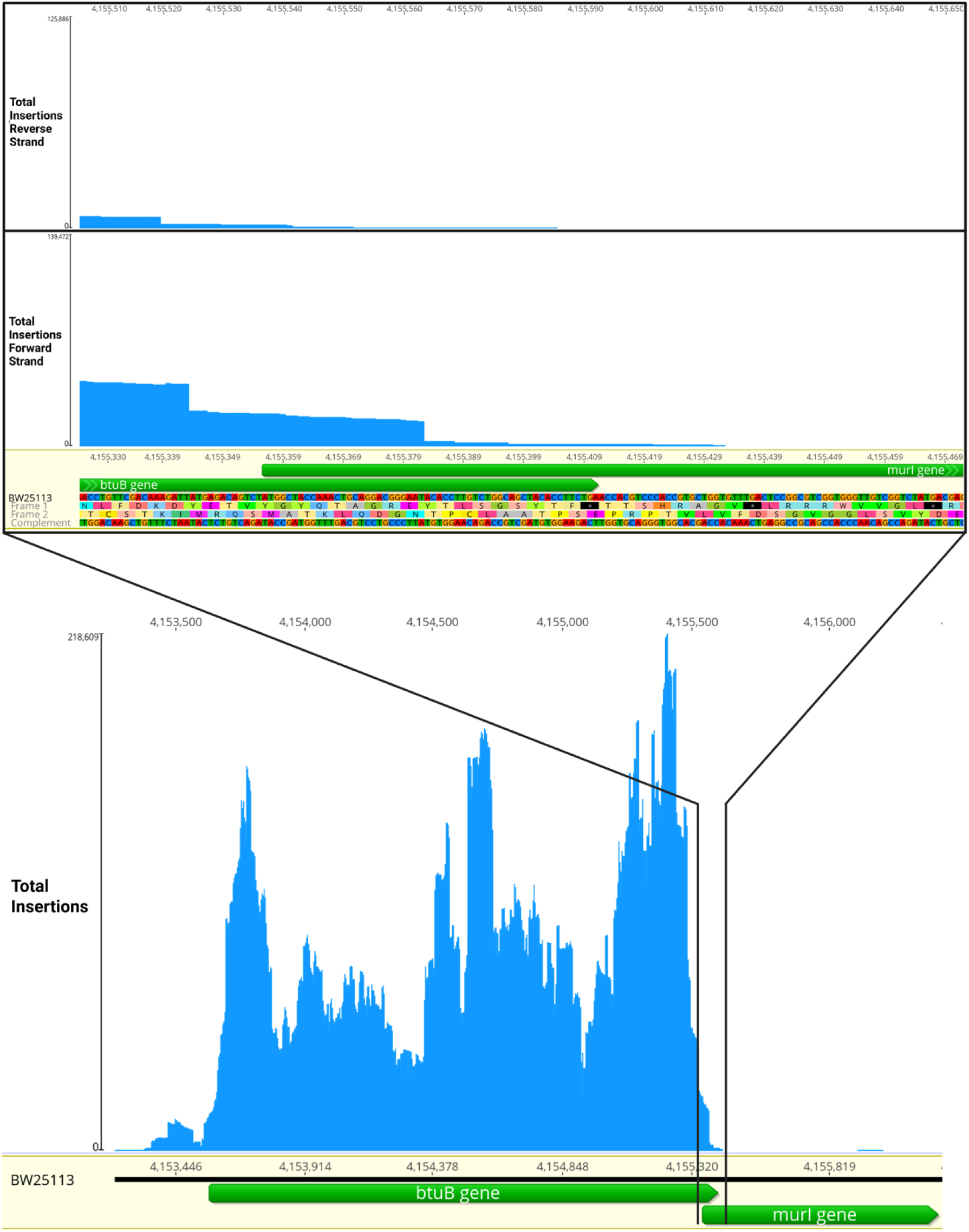
transposon insertion frequency in the *btuB/murI* region. The overlap region at the 3’ end of the *btuB* gene is shown expanded above, with the directions of each insertion mapped; upper insert panel: insertions in the reverse orientation, lower insert panel: insertions in the forward direction.

**Supplementary table 1:**
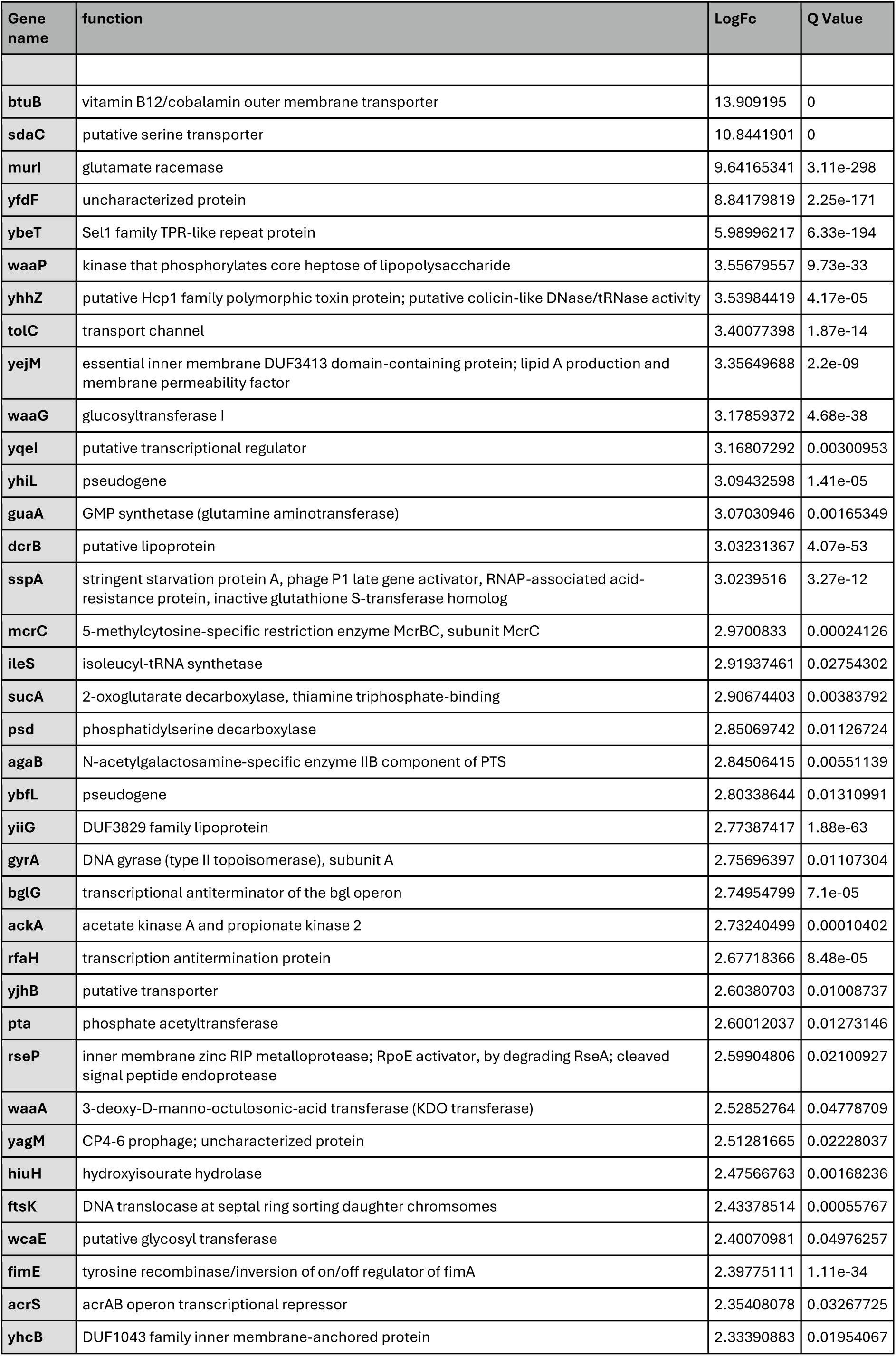
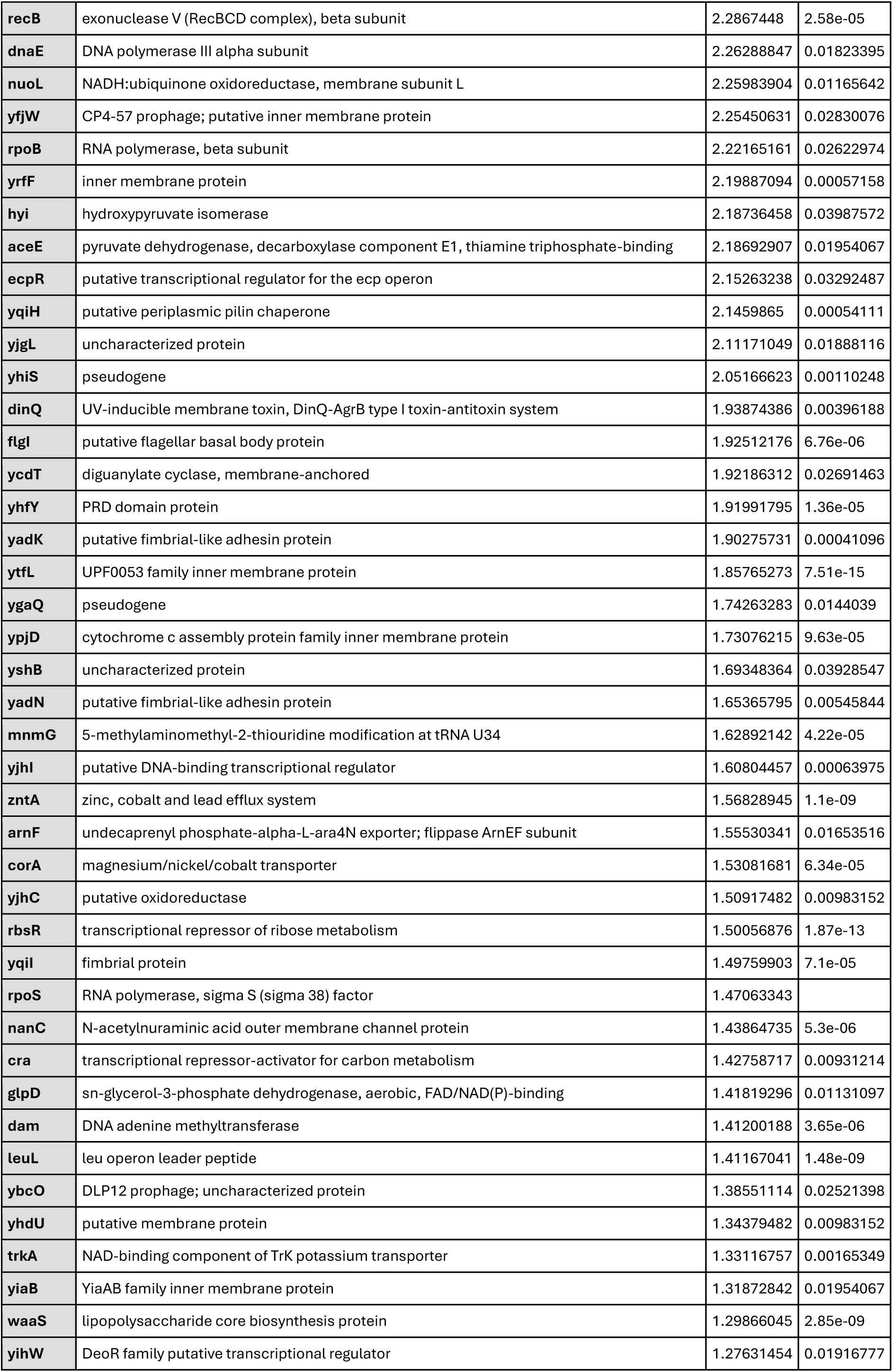
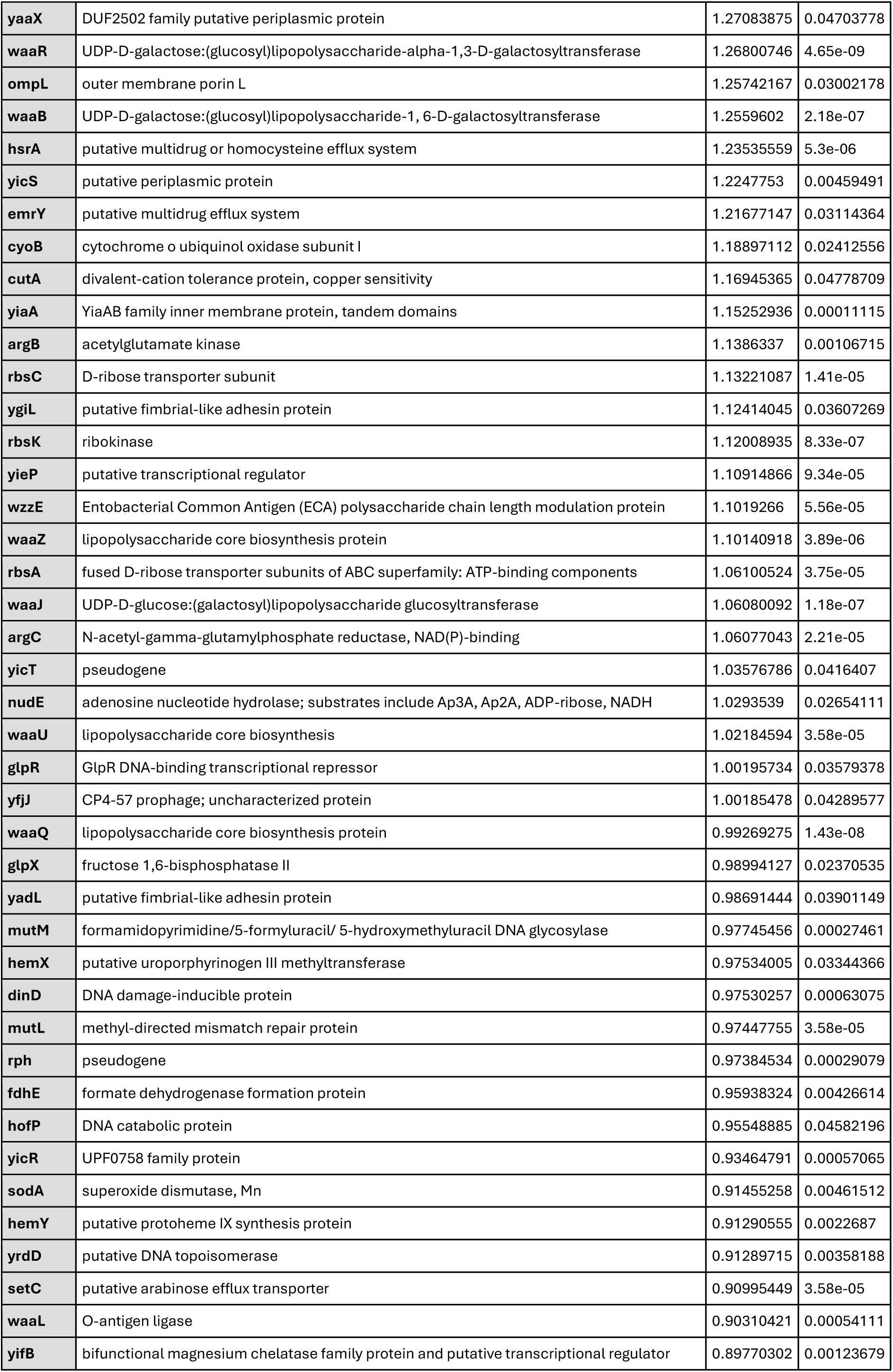
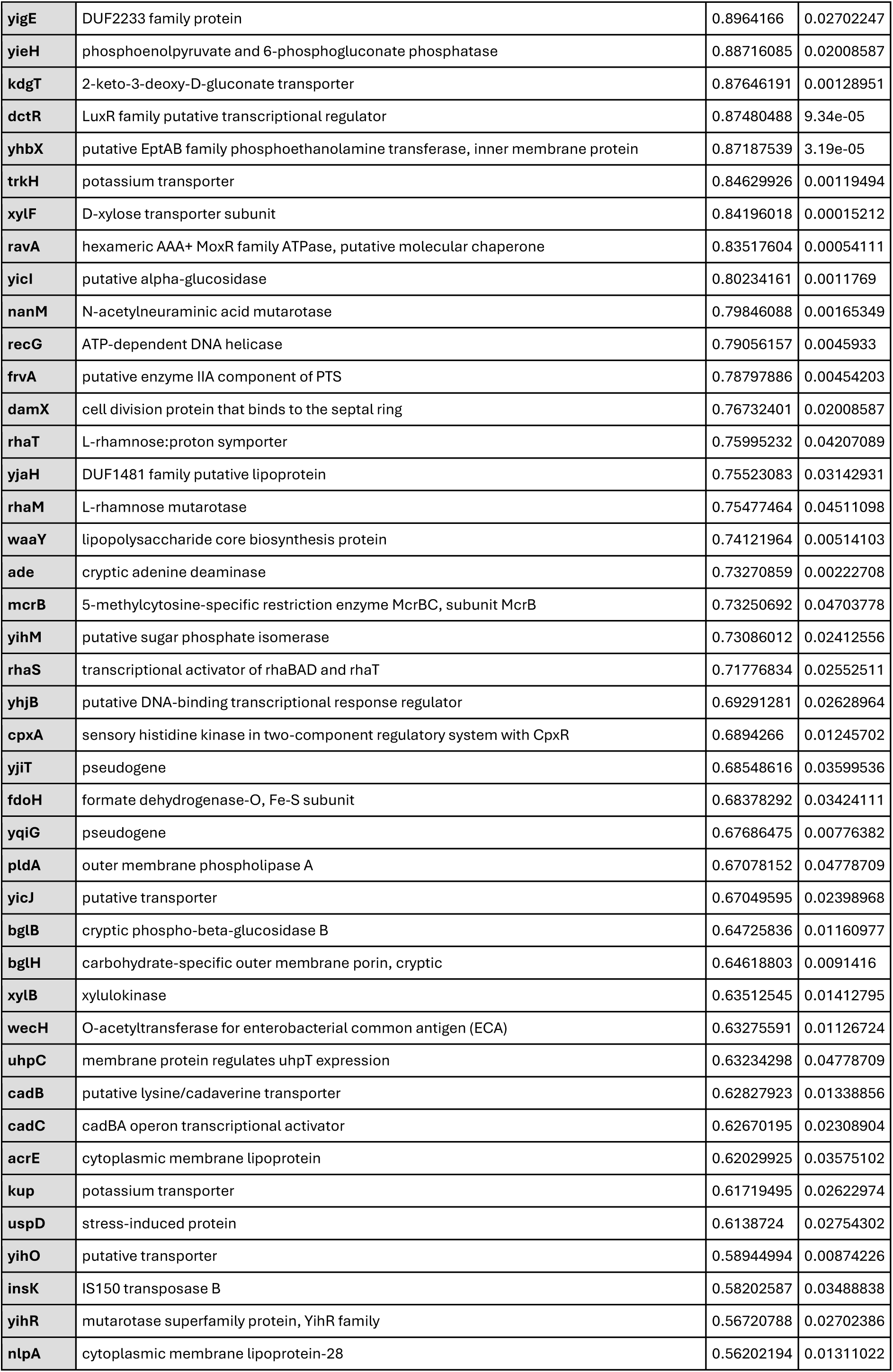
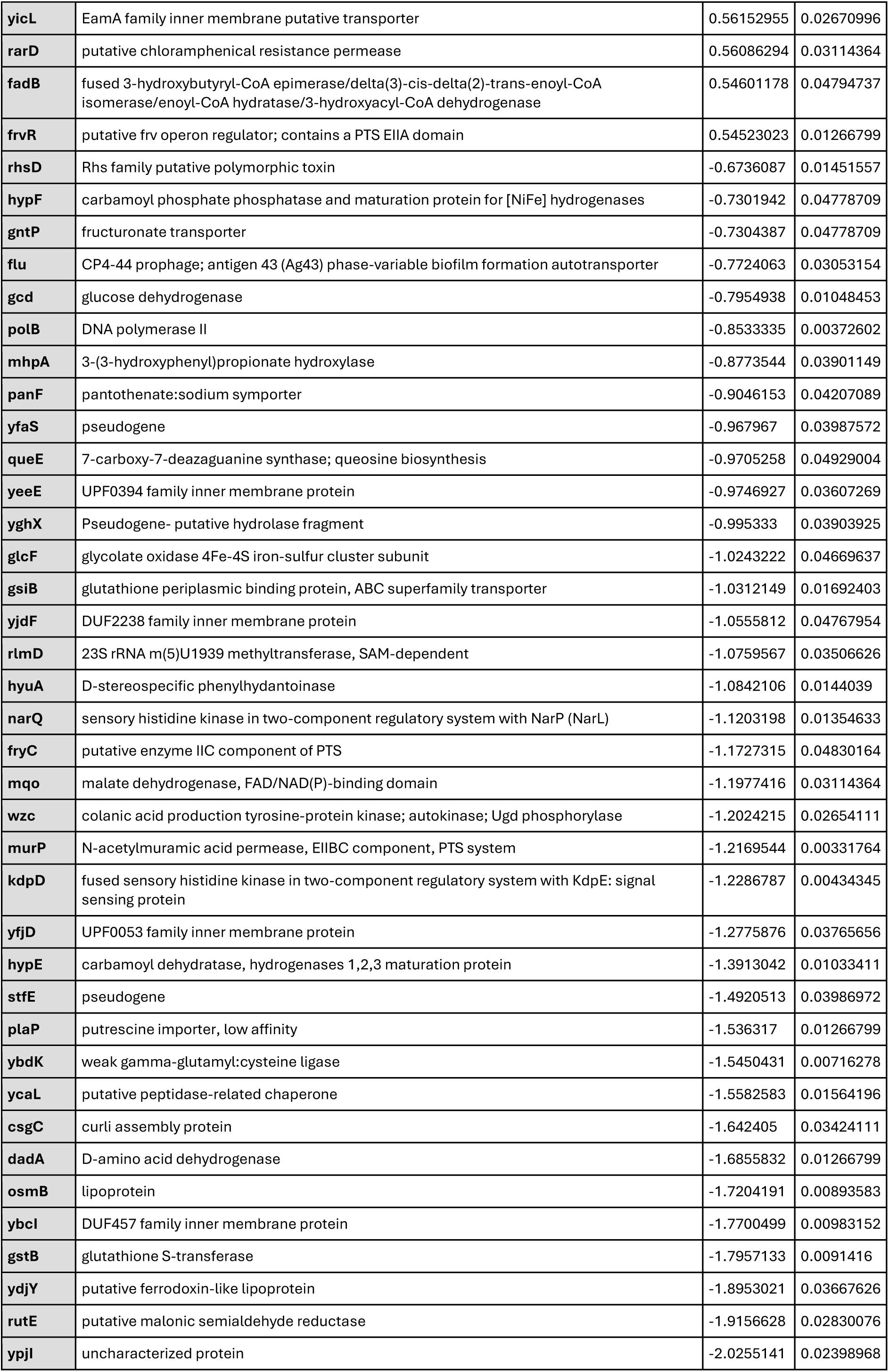
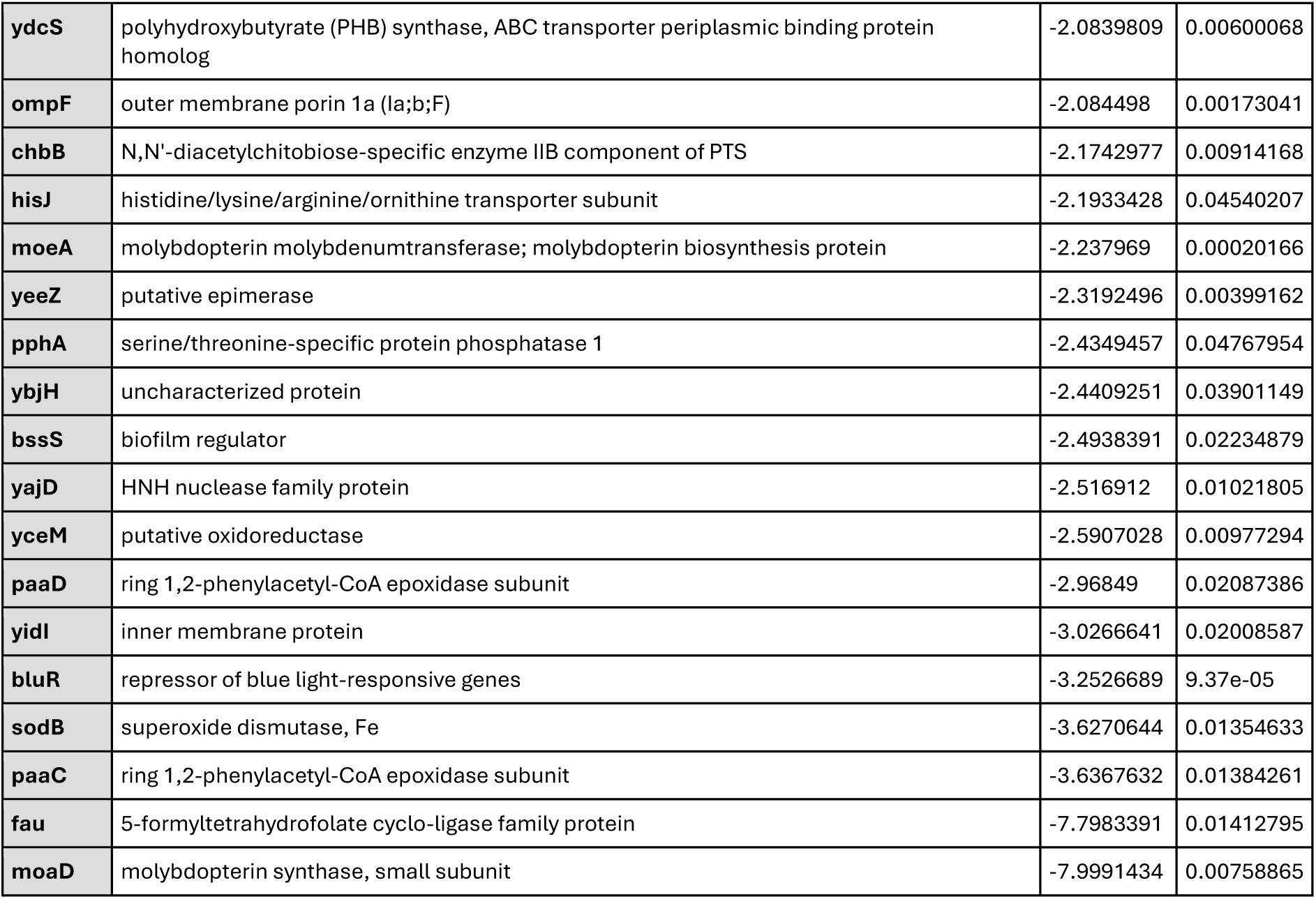
list of all significantly (Q < 0.05) over- or under-represented genes from TraDIS sequencing following Arnold challenge compared to the control input library. The gene name, predicted function, Log_10_FC and Q values are shown. Positive Log_2_FC represent mutants in genes that showed better survival under phage challenge, whereas negative Log_2_FC represent genes that showed poorer survival after phage challenge relative to the input pool. Note that the top two genes *btuB* and *sdaC* had Q values <10^-300^, so the TraDIS pipeline annotated these as 0.

## References

Alquethamy, S. F., Adams, F. G., Maharjan, R., Delgado, N. N., Zang, M., Ganio, K., Paton, J. C., Hassan, K. A., Paulsen, I. T., McDevitt, C. A., Cain, A. K., & Eijkelkamp, B. A. (2021). The Molecular Basis of Acinetobacter baumannii Cadmium Toxicity and Resistance. Appl Environ Microbiol, 87(22), e0171821. 10.1128/AEM.01718-21

Antimicrobial Resistance, C. (2022). Global burden of bacterial antimicrobial resistance in 2019: a systematic analysis. Lancet, 399(10325), 629–655. 10.1016/S0140-6736(21)02724-0

Baba, T., Ara, T., Hasegawa, M., Takai, Y., Okumura, Y., Baba, M., Datsenko, K. A., Tomita, M., Wanner, B. L., & Mori, H. (2006). Construction of *Escherichia coli* K-12 in-frame, single-gene knockout mutants: the Keio collection. Molecular Systems Biology, 2(1), 2006.0008. 10.1038/msb4100050

Barquist, L., Mayho, M., Cummins, C., Cain, A. K., Boinett, C. J., Page, A. J., Langridge, G. C., Quail, M. A., Keane, J. A., & Parkhill, J. (2016). The TraDIS toolkit: sequencing and analysis for dense transposon mutant libraries. Bioinformatics, 32(7), 1109–1111. 10.1093/bioinformatics/btw022

Bouras, G., Nepal, R., Houtak, G., Psaltis, A. J., Wormald, P.-J., & Vreugde, S. (2022). Pharokka: a fast scalable bacteriophage annotation tool. Bioinformatics, 39(1). 10.1093/bioinformatics/btac776

Christen, B., Abeliuk, E., Collier, J. M., Kalogeraki, V. S., Passarelli, B., Coller, J. A., Fero, M. J., McAdams, H. H., & Shapiro, L. (2011). The essential genome of a bacterium. Mol Syst Biol, 7, 528. 10.1038/msb.2011.58

Cooper, M. A., & Shlaes, D. (2011). Fix the antibiotics pipeline. Nature, 472(7341), 32. 10.1038/472032a

Cowley, L. A., Low, A. S., Pickard, D., Boinett, C. J., Dallman, T. J., Day, M., Perry, N., Gally, D. L., Parkhill, J., Jenkins, C., & Cain, A. K. (2018). Transposon Insertion Sequencing Elucidates Novel Gene Involvement in Susceptibility and Resistance to Phages T4 and T7 in Escherichia coli O157. mBio, 9(4). 10.1128/mBio.00705-18

Cumby, N., Edwards, A. M., Davidson, A. R., & Maxwell, K. L. (2012). The bacteriophage HK97 gp15 moron element encodes a novel superinfection exclusion protein. J Bacteriol, 194(18), 5012–5019. 10.1128/JB.00843-12

Cumby, N., Reimer, K., Mengin-Lecreulx, D., Davidson, A. R., & Maxwell, K. L. (2015). The phage tail tape measure protein, an inner membrane protein and a periplasmic chaperone play connected roles in the genome injection process of E. coli phage HK97. Mol Microbiol, 96(3), 437–447. 10.1111/mmi.12918

Datsenko, K. A., & Wanner, B. L. (2000). One-step inactivation of chromosomal genes in Escherichia coli K-12 using PCR products. Proc Natl Acad Sci U S A, 97(12), 6640–6645. 10.1073/pnas.120163297

Domka, J., Lee, J., & Wood, T. K. (2006). YliH (BssR) and YceP (BssS) Regulate *Escherichia coli* K-12 Biofilm Formation by Influencing Cell Signaling. Applied and Environmental Microbiology, 72(4), 2449–2459. doi:10.1128/AEM.72.4.2449-2459.2006

Gilbey, T., Ho, J., Cooley, L. A., Petrovic Fabijan, A., & Iredell, J. R. (2019). Adjunctive bacteriophage therapy for prosthetic valve endocarditis due to Staphylococcus aureus. Med J Aust, 211(3), 142–143 e141. 10.5694/mja2.50274

Goodall, E. C. A., Robinson, A., Johnston, I. G., Jabbari, S., Turner, K. A., Cunningham, A. F., Lund, P. A., Cole, J. A., & Henderson, I. R. (2018). The Essential Genome of Escherichia coli K-12. mBio, 9(1). 10.1128/mBio.02096-17

Harding, K. R., Malone, L. M., Kyte, N. A. P., Jackson, S. A., Smith, L. M., & Fineran, P. C. (2025). Genome-wide identification of bacterial genes contributing to nucleus-forming jumbo phage infection. Nucleic Acids Res, 53(3). 10.1093/nar/gkae1194

Langmead, B., & Salzberg, S. L. (2012). Fast gapped-read alignment with Bowtie 2. Nat Methods, 9(4), 357–359. 10.1038/nmeth.1923

Letellier, L., Plancon, L., Bonhivers, M., & Boulanger, P. (1999). Phage DNA transport across membranes. Res Microbiol, 150(8), 499–505. 10.1016/s0923-2508(99)00107-2

Likhacheva, N. A., Khrenova, E. A., & Sineokii, S. P. (1990). [Genetic control of resistance of Escherichia coli K12 to phage C1]. Mol Gen Mikrobiol Virusol(12), 12–15. https://www.ncbi.nlm.nih.gov/pubmed/2150690 (Geneticheskii kontrol’ rezistentnosti Escherichia coli K12 k fagu C1.)

Likhacheva, N. A., Samsonov, V. V., Samsonov, V. V., & Sineoky, S. P. (1996). Genetic control of the resistance to phage C1 of Escherichia coli K-12. J Bacteriol, 178(17), 5309–5315. 10.1128/jb.178.17.5309-5315.1996

Lindberg, A. A. (1973). Bacteriophage receptors. Annu Rev Microbiol, 27, 205–241. 10.1146/annurev.mi.27.100173.001225

Maher, C., Maharjan, R., Sullivan, G., Cain Amy, K., & Hassan Karl, A. (2022). Breaching the Barrier: Genome-Wide Investigation into the Role of a Primary Amine in Promoting E. coli Outer-Membrane Passage and Growth Inhibition by Ampicillin. Microbiology Spectrum, 10(6), e03593–03522. 10.1128/spectrum.03593-22

O’Neill, J. (2014). ‘Review on Antimicrobial Resistance. Antimicrobial Resistance: Tackling a Crisis for the Health and Wealth of Nations. 2014 (HM Government, UK., Issue.

Salmond, G. P., & Fineran, P. C. (2015). A century of the phage: past, present and future. Nat Rev Microbiol, 13(12), 777–786. 10.1038/nrmicro3564

Samsonov, V. V., Samsonov, V. V., & Sineoky, S. P. (2002). DcrA and dcrB Escherichia coli genes can control DNA injection by phages specific for BtuB and FhuA receptors. Res Microbiol, 153(10), 639–646. 10.1016/s0923-2508(02)01375-x

Seeman, T. (2020). Shovill. *Available at:* https://github.com/tseemann/shovill.

Washizaki, A., Yonesaki, T., & Otsuka, Y. (2016). Characterization of the interactions between Escherichia coli receptors, LPS and OmpC, and bacteriophage T4 long tail fibers. Microbiologyopen, 5(6), 1003–1015. 10.1002/mbo3.384

